# Dissecting limiting factors of the Protein synthesis Using Recombinant Elements (PURE) system

**DOI:** 10.1101/099838

**Authors:** Jun Li, Chi Zhang, Poyi Huang, Erkin Kuru, Eliot T. C. Forster-Benson, Taibo Li, George M. Church

**Affiliations:** Department of Genetics, Harvard Medical School, 77 Avenue Louis Pasteur, Boston, MA 02115, USA.; Wyss Harvard Institute of Biologically Inspired Engineering, 3 Blackfan Circle, Boston, MA 02115, USA.; Ravenwood High School, 1724 Wilson Pike, Brentwood, TN 37027, USA.; Department of Electrical Engineering and Computer Science, Massachusetts of Institute of Technology, 77 Massachusetts Avenue, Cambridge, MA 02139, USA.

**Author notes:** Correspondences should be addressed to Jun Li at or George M. Church at.

## Abstract

Reconstituted cell-free protein synthesis systems such as the Protein synthesis Using Recombinant Elements (PURE) system give high-throughput and controlled access to *in vitro* protein synthesis. Here we show that compared to the commercial S30 crude extract based RTS 100 *E. coli* HY system, the PURE system has less mRNA degradation and produces ~4-fold more total protein. However the majority of these polypeptides are partially translated or inactive since the signal from firefly luciferase (Fluc) translated in PURE is only ~2/3^rd^ of that measured using the S30 crude extract system. Both of the two systems suffer from low ribosome recycling efficiency when translating proteins from 90 kD to 220 kD. A systematic fed-batch analysis of PURE shows replenishment of 6 small molecule substrates individually or in combination prior to energy depletion increased Fluc protein yield by ~1.5 to ~2-fold, while accumulation of inorganic phosphate contributes to reaction termination. Additionally, while adding EF-P to PURE reduced total protein translated, it also increased the fraction of active product and reduced partial translated product probably by slowing down the translation process. Finally, ArfA, rather than YaeJ or PrfH, helped reduce ribosome stalling when translating Fluc and improved system productivity in a template-dependent fashion.

## INTRODUCTION

Cell-free protein synthesis (CFPS) enables rapid expression of functional proteins from their genes. It is well applied in display technologies^1, 2^, unnatural amino acids incorporation into proteins^3, 4^, and *in vitro* protein evolution^5^ and provides a platform to study protein translation and folding. Conventional CFPS systems use cell lysates from *E. coli* cells^6^, wheat germ^7^ or rabbit reticulocytes^8^ to supply the protein translation machinery and a recombinant T7 RNA polymerase to couple transcription to translation. As a more stripped-down and defined CFPS system, “Protein synthesis Using Recombinant Elements” (PURE) system is reconstituted solely from elements essential to *E. coli* translation^9^. For example, as opposed to the *E. coli* extract-based S30 system^10^, the PURE system is avoid of cellular contaminants such as proteases and nucleases inhibitory to protein synthesis^11^.

Conventional batch format PURE experiment has the capacity for a yield of over 100 μg/ml^12^, while batch format S30 system, such as RTS 100 *E. coli* HY system, claims a yield range from 40-400 μg/ml^13^. However, the batch PURE reaction often lasts for a shorter period of time than the batch format S30 system^14, 15^. Depletion of primary and secondary energy substrate^16, 17^, accumulation of inhibitory byproducts^18, 19^, and shortage of nutrients such as tRNAs, amino acids and other small molecule substrates^20–22^ could be possible explanations for this significant difference. Also, the higher translation durations of S30 system is suggested to involve presence of additional unidentified cellular factors capable of enhancing protein translation and folding^23^. For example, supplementing the PURE system with fractions from a size-exclusion separation of the S30 extract significantly increases the yield and activity to a level close to that of the S30 system^24^. Although the exact composition of these PURE-enhancing fractions are still not clear, it likely contains at least a chaperone as adding the GroEL/ES chaperone system to the PURE system alone can increase the functional firefly luciferase yield by 60%^25^.

Additionally, the PURE system produces a significant amount of partially translated products most likely due to ribosome stalling on mRNA^25, 26^. In *E. coli,* ribosome rescue is accomplished through three systems: 1-trans-translation mediated by transfer messenger RNA (tmRNA) and a partner protein, SmpB^27^, 2-ArfA with **<u>r</u>**elease **<u>f</u>**actor-**<u>2</u>** (RF2)^28, 29^ which can release stalled ribosomes from mRNAs with no in-frame stop codons and 3-YaeJ (ArfB) ^30^. In addition, the **<u>e</u>**longation **<u>f</u>**actor **<u>P</u>** (EF-P) alleviates ribosome stalling by enhancing translocation of proline codon stretches^31, 32^ and **<u>e</u>**longation **<u>f</u>**actor **<u>4</u>** (EF4) stimulates protein synthesis under high Mg^2+^ concentration^33^. This list is still growing. For example, PrfH, a pseudogene from *E. coli K12*, which contains a highly conserved GGQ motif associated with peptidyl hydrolysis activity, has been proposed to respond to signals other than conventional stop codons^34^. However, no experimental evidence has been reported on the function of PrfH.

Here, we performed a side by side comparison between the PURE and RTS 100 *E. coli* HY S30 systems in terms of mRNA stabilities, total, active and partially translated product yields and ribosome recycling efficiencies. To test and overcome the effects of substrate depletion, we supplemented the initial reaction with higher concentrations of tRNAs, amino acids (AA), the reducing agent dithiothreitol (DTT), 10-Formyl-5,6,7,8-tetrahydrofolic acid (10-CH0-THF), NTPs, the secondary energy substrate creatine phosphate and Mg^2+^ individually or in combination to the PURE system. In parallel, we monitored the accumulation of the inhibitory byproduct, inorganic phosphate, using a chemical assay. Besides, we supplemented the PURE system with purified missing protein factors present in cells and S30 but absent in PURE to study how these factors affect translation. We show adding EF-P increased the active product yield while decreased total and partially translated product yields even with a template with no poly-proline stretches, possibly by slowing down translation initiation or elongation. EF4 increased active, total and partial translated products in PURE with elevated Mg^2+^ concentrations. ArfA, but not YaeJ or PrfH, helped ribosomes stalled at templates with no stop codons release when large protein was expressed in the PURE system.

## RESULTS

### Comparing PURE system transcription and translation with RTS 100 *E. coli* HY S30 system

The PURE system is reconstituted from T7 RNA polymerase, ribosomes, initiation factors (IF1, IF2, and IF3), elongation factors (EF-G, EF-Tu, and EF-Ts), release factors (RF1, RF2, and RF3), ribosome recycling factor (RRF), 20 aminoacyl-tRNA synthetases (ARSs), methionyl-tRNA transformylase (MTF), 10-CH0-THF, 46 tRNAs, 20 amino acids, NTPs, DTT, creatine phosphate, creatine kinase, myokinase, nucleoside-diphosphate kinase, and pyrophosphatase. To begin, we assessed the duration of PURE protein synthesis with three plasmid templates encoding proteins with different lengths: Namely firefly luciferase with an N-terminal tetracysteine tag (TC-Fluc, 63 kD), a fusion of HaloTag, mCherry and TolA (HT-Ch-TolA, 90 kD) and a fusion of HaloTag, EntF, mCherry and TolA (HT-EntF-Ch-TolA, 220 kD). TolA is a C-terminal 171-amino-acid alpha-helical spacer excised from *E. coli* TolA domain II. EntF is a multidomain enzyme from *E. coli* enterobactin biosynthetic pathway.

We first measured active TC-Fluc produced in the PURE system over time with a luciferase assay by converting relative luminescence units to the actual amount of TC-Fluc via a premeasured standard curve. The amount of active TC-Fluc produced increased as the reaction went on and reached a maximum of 114.3 ± 3.4 nmol/L at 90 min (Supplementary Figure 1A). We also tested the expression of HT-Ch-TolA and HT-EntF-Ch-TolA. We observed a protein-specific, unique fluorescence pattern for both HT-Ch-TolA and HT-EntF-Ch-TolA (Supplementary Figure 1B). This strongly suggests that besides full length products, partially translated polypeptides were also abundant, possibly due to premature termination or pausing during translation^26, 35^. Finally, the intensity of bands of full length and partial products reached a maximum ~ 2 h (Supplementary Figure 1B), indicative of the stopping point for the PURE reaction under these conditions.

In order to compare the transcription and translation between the PURE system and the S30 system, we chose TC-Fluc as a model protein. Circular plasmid encoding TC-Fluc was added to PURE system and S30 system to the same final concentration of 10 ng/μl. The PURE system reaction was incubated at 37°C for 2 h, while the S30 system reaction was incubated at 30°C for 6 h. Analysis of total RNA extracted from the PURE system reaction revealed 23S, 16S, 5S ribosomal RNA (rRNA), tRNAs and the 1.9 kb T7 TC-Fluc mRNA when compared to the native 23S, 16S, 5S rRNA and tRNAs as markers (Figure 1A-B). Bands larger than the 2.9 kb 23S rRNA (indicated by asterisk *) also appeared on the gel, probably due to the inefficient termination of T7 polymerase^36^. No degraded mRNA template was found in the PURE reaction, suggesting that the poorer sustainability of the PURE system is not incurred by ribosome stalling due to truncated mRNA templates (Figure 1A). In contrast, the total RNA from S30 system reaction contained not only T7 TC-Fluc mRNA appearing at approximately 1.9 kb, but also a smear of different bands smaller than the 16S band indicating partial degradation of the RNA template (Figure 1B).

**Figure 1.**
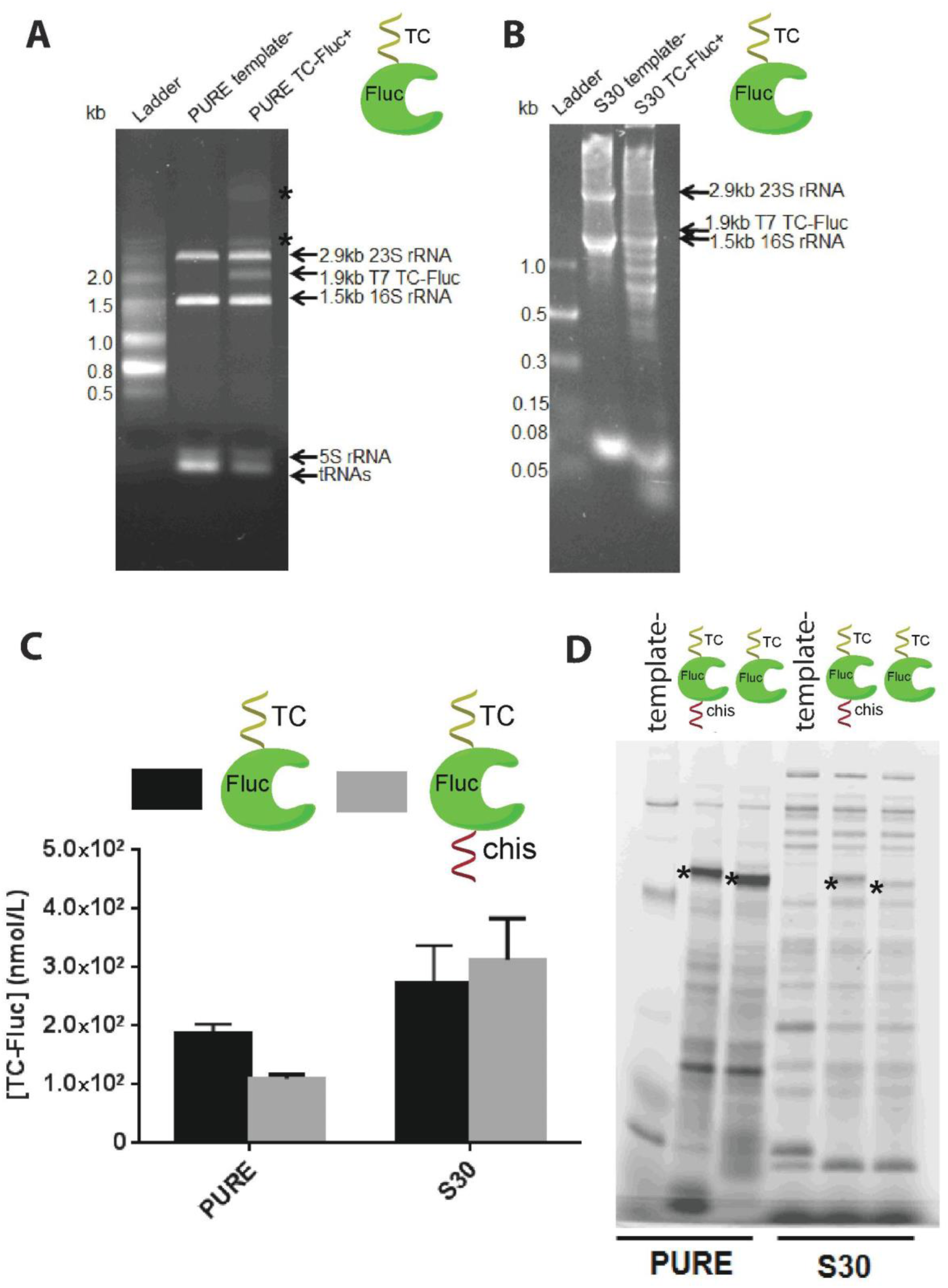
Comparison of transcription and translation between PURE system and RTS 100 *E. coli* HY S30 cell extract system. **A.** RNA denaturing gel of PURE reaction transcribing TC-Fluc mRNA. PURE system with no plasmid template and plasmid encoding TC-Fluc were incubated at 37°C for 2 h. Total RNA were purified by phenol/chloroform extraction and isopropanol precipitation from equal volumes of the two reactions and then run on lane 2 and 3, respectively. **B.** RNA denaturing gel of S30 extract transcribing TC-Fluc mRNA. S30 extract with no plasmid template and plasmid encoding TC-Fluc were incubated at 30°C for 6 h. Total RNAs were purified by phenol/chloroform extraction and isopropanol precipitation from equal volumes of the two reactions and then run on lane 2 and 3, respectively. **C.** Assessment of active TC-Fluc and TC-Fluc-chis proteins translated in PURE system and S30 extract. Equal volume aliquots were taken for luciferase assay to measure the actual amount of active TC-Fluc produced. Values represent averages and error bars are ± standard deviations, with n=3. **D.** Assessment of total TC-Fluc and TC-Fluc-chis produced in PURE system and S30 extract. TC-Fluc and TC-Fluc-chis synthesized in **C** were incubated with FlAsH-EDT_2_ biarsenical labeling reagent and analyzed on a SDS-PAGE. The gel was scanned by a typhoon scanner with filter set (508nmE×/528nmEm). Asterisk^*^ indicates full lenth product.

To compare the efficiency of translation, TC-Fluc and TC-Fluc with a C-terminal His tag (TC-Fluc-chis) were synthesized in PURE and S30 system and the yields were quantified by luciferase assays in equal volumes of PURE and S30 system. Compared to the PURE system, S30 system produced ~1.5-fold and ~3-fold more functional TC-Fluc and TC-Fluc-chis, respectively (Figure 1C). We also quantified yields of full-length proteins, by mixing equal volume of the PURE and S30 system reactions with FlAsH-EDT_2_, a biarsenical labeling reagent to label proteins with a TC tag, and running the samples on a SDS-PAGE and detecting fluorescence. Although the control reactions with no plasmid template showed some fluorescent bands and smearing indicative of non-specific FlAsH-EDT_2_ labeling (lane 1 and 4 in Figure 1D), the full length and incomplete TC-Fluc and TC-Fluc-chis could still be clearly detected in both of the two systems (Figure 1D). Based on the band intensity, the PURE system produced ~4-fold more full length TC-Fluc and TC-Fluc-chis compared to the S30 system. The significantly lower activity of TC-Fluc and TC-Fluc-chis in PURE (Figure 1C) suggests that a large portion of the proteins produced by the PURE system is not active, most likely due to protein misfolding.

Majority of the bands and smearing other than the full length product in S30 system came from non-specific labelling of FlAsH-EDT_2_. Since the PURE system produced also a large amount of incomplete products of different lengths, we wanted to minimize the interference from non-specific labeling. Therefore, we expressed three HaloTag fusion proteins HT-Ch-TolA, HT-EntE-Ch-TolA (EntE is also a multidomain enzyme from *E. coli* enterobactin biosynthetic pathway) and HT-EntF-Ch-TolA in the PURE and S30 systems. HT-Ch-TolA, HT-EntE-Ch-TolA and HT-EntF-Ch-TolA were prepared as linear DNA templates with two versions: with and without stop codon. We labeled the products with HaloTag TMR ligand, run them on SDS-PAGE and compared the in-gel fluorescence. The PURE system produced ~2-fold more full length HT-Ch-TolA, HT-EntE-Ch-TolA and HT-EntF-Ch-TolA compared to the S30 system (Figure 2). Consistent with the TC labeling, HT labeling confirmed that the PURE system produced larger amounts of incomplete polypeptide products than S30 for all of the three proteins (Figure 2), each with characteristic unique patterns. This suggests that PURE system might be lacking functionalities against premature protein termination perhaps due to ribosome stalling. Templates with and without stop codon didn’t show much difference on the yield of full length and partially translated products for all of the three proteins in both PURE and S30 systems (Figure 2), indicating low ribosome recycling efficiency.

**Figure 2.**
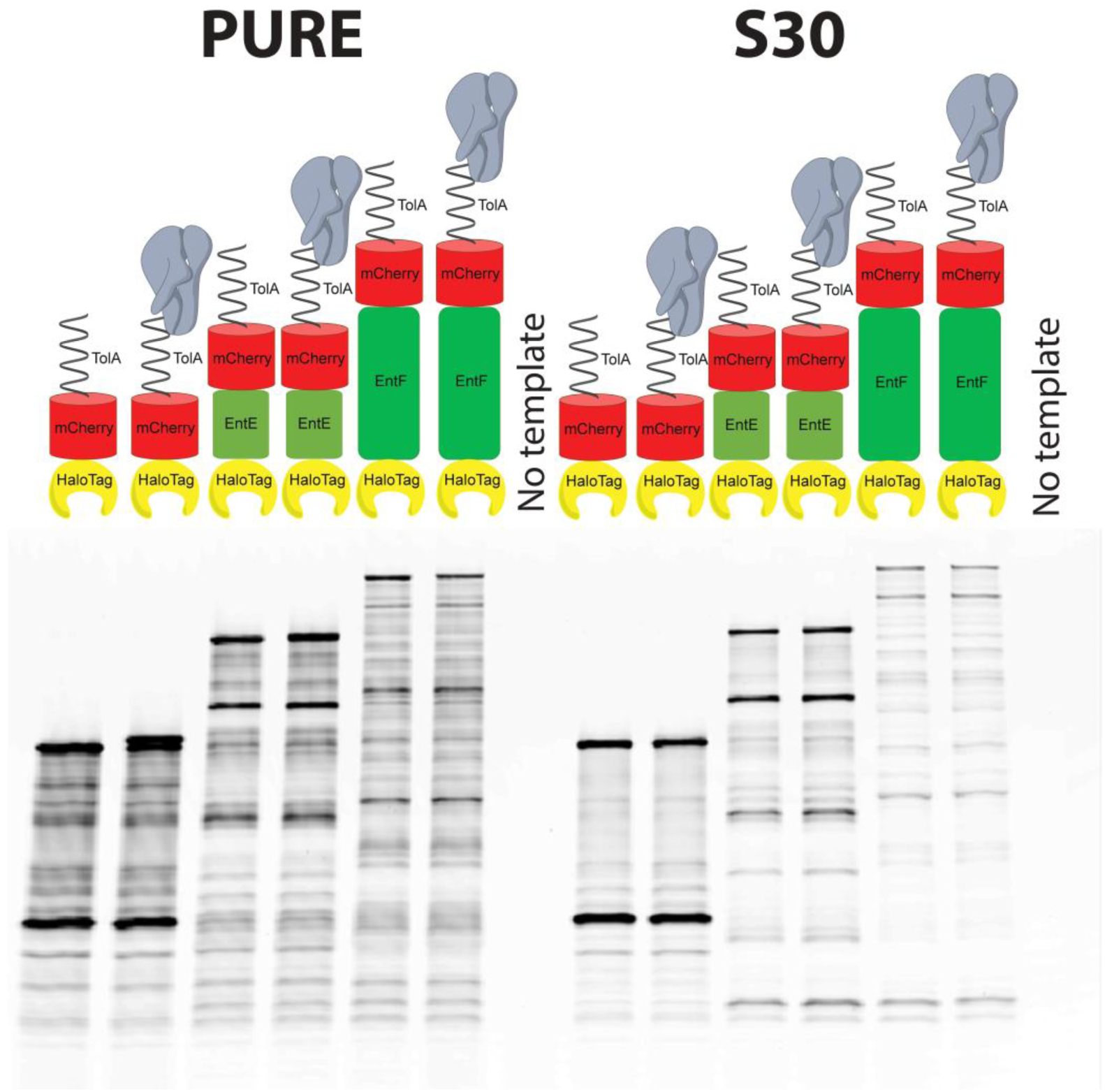
Assessment of PURE and RTS 100 *E. coli* HY S30 system translation by production of HaloTag fusion proteins using linear DNA templates. HT-Ch-TolA, HT-EntE-Ch-TolA and HT-EntF-Ch-TolA with and without stop codon (90 kD, 160 kD, 220 kD) were synthesized in the PURE system and S30 extract and then incubated with HaloTag TMR Ligand. Equal volume aliquots of samples from PURE system and S30 extract reactions were analyzed on a SDS-PAGE and then scanned by a typhoon scanner with filter set (555nmE×/580nmEm). Lane 1, 3, 5, 8, 10, 12 are HaloTag fusion proteins with stop codon. Lane 2, 4, 6, 9, 11, 13 are HaloTag fusion proteins without stop codon. Lane 1 and 14 are negative controls with no template. (HT: HaloTag; Ch: mCherry. TolA is a C-terminal 171-amino-acid alpha-helical spacer excised from E. coli TolA domain II. EntE and EntF are multidomain enzymes from E. coli enterobactin biosynthetic pathway.)

### Substrate replenishment with fed-batch PURE system

Previously we showed that optimization of IFs, EFs, RFs and RRF concentrations can increase the yields of active Fluc in the PURE system to various extents^25^. Historically, nonproductive energy consumption, secondary energy source depletion, and amino acid exhaustion have been identified as the primary reasons for early termination of CFPS systems that use plasmid DNA templates^37, 38^. In order to study whether small molecule substrate availability restricted the productivity of the PURE system, we performed fed-batch experiments by replenishing tRNAs, AA, DTT, 10-CH0-THF, NTPs, the secondary energy substrate creatine phosphate and Mg^2+^ individually or in combination at 15 min, 30 min, 45 min, 60 min or 75 min after the reaction started and measured the amount of active TC-Fluc produced at 90 min, the time-point at which the TC-Fluc synthesis stopped (Supplementary Figure 1A).

Feeding 0.3 mM AA or 27 0D_260_/ml tRNAs once at different time points only slightly increased active TC-Fluc yield (Figure 3A-B). But when combined together, it increased the TC-Fluc signal maximum ~80% above that of the control (water supplement only) when fed at 75 min (Figure 3C). The system uses DTT as reducing agent to prevent intramolecular and intermolecular disulfide bonds from forming between cysteine residues of proteins. As DTT is not stable at 37°C, we replenished 1 mM DTT to the system and found that feeding at 15 min, 30 min, 45 min, 60 min only slightly improved the yield of active TC-Fluc, while feeding at 75 min gave an increase of ~50% compared to the control (Figure 3D), indicating that DTT might start to degrade at about 75 min after the reaction started. In the PURE system, the enzyme MTF formylates Met-tRNA^fMet^ using 10-CH0-THF to form fMet-tRNA^fMet^ for translation initiation. Replenishing 10 μg/ml 10-CH0-THF boosted active TC-Fluc yield a maximum of ~50% when fed at 60 min or 75 min (Figure 3E).

**Figure 3.**
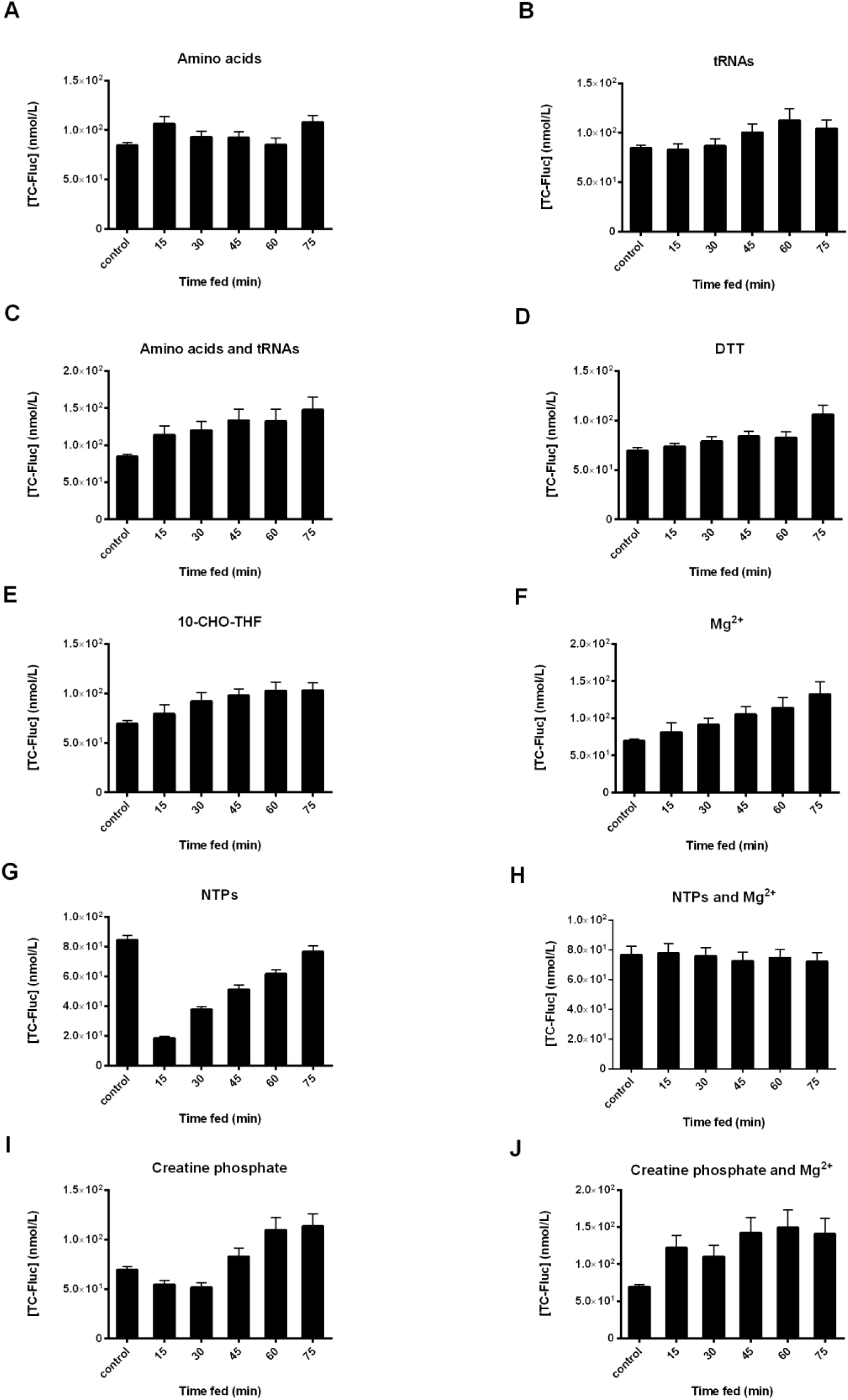
Replenishing small molecule substrates in the PURE system protein synthesis at different time points. PURE system reactions synthesizing TC-Fluc were set up and incubated at 37°C. Small molecule substrates as indicated were fed to the reaction at 15 min, 30 min, 45 min, 60 min or 75 min. Control reactions were fed equivalent volumes of water at 0 min. Overall functional TC-Fluc produced was assessed by luciferase assay at 90 min. Values represent averages and error bars are ± standard deviations, with n=3. PURE system feed supplements of **A.** 0.3 mM amino acids; **B.** 27 OD_260_/ml tRNAs; **C.** 0.3 mM amino acids and 27 OD_260_/ml tRNAs in combination; **D.** 1 mM DTT; **E.** 10 μg/ml 10-CHO-THF; **F.** 2 mM magnesium acetate; **G.** 1.6 mM ATP, 1.6 mM GTP, 0.8 mM CTP and 0.8 mM UTP; **H.** 4.8 mM magnesium acetate together with 1.6 mM ATP, 1.6 mM GTP, 0.8 mM CTP and 0.8 mM UTP; **I.** 20 mM creatine; and **J.** 20 mM creatine phosphate and 2 mM magnesium acetate in combination.

Not surprisingly, the depletion of creatine phosphate as the secondary energy source was coupled with a rapid increase in inorganic phosphate which sequesters magnesium present in the system necessary for operation of the translation apparatus. We observed that inorganic phosphate concentrations reached ~20 mM after incubating the reaction at 37°C for 3 h (Supplementary Figure 2). Thus we fed 2 mM magnesium acetate to offset the deleterious effects of magnesium sequestration and observed a maximum of ~50% increase in active TC-Fluc yield when feeding at 75 min (Figure 3F). Feeding 4.8 mM magnesium acetate had an inhibitory effect on TC-Fluc translation (Supplementary Figure 3A). Feeding 1.6 mM ATP, 1.6 mM GTP, 0.8 mM CTP and 0.8 mM UTP inhibited translation of TC-Fluc (Figure 3G) regardless of feeding time points, probably due to NTPs’ chelation with magnesium at a 1:1 molar ratio. Consistently, feeding NTPs at later time points had less inhibitory effect than feeding at earlier time points, that could be explained by NTPs being consumed as the reaction went on (Figure 3G). To offset this chelating effect, we also supplemented 4.8 mM magnesium acetate together with 1.6 mM ATP, 1.6 mM GTP, 0.8 mM CTP and 0.8 mM UTP to the system and observed no effect on active TC-Fluc yield, suggesting that the availability of NTPs is not a key limiting factor for TC-Fluc synthesis in the PURE system (Figure 3H). Creatine phosphate was fed at its initial concentration of 20 mM and reached maximum increase in active TC-Fluc yield of ~60% when fed at 75 min (Figure 3I). When creatine phosphate is combined with 2 mM magnesium acetate, the yield increased to ~2-fold of that of the control at maximum (Figure 3J), although higher magnesium acetate concentrations (4.8 mM) did not significantly change the yield observed (Supplementary Figure 3B).

Based on these observations on the effects of individual components and time points, we replenished the reaction at the feeding time 75 min with a combination of AA, tRNAs, DTT, 10-CH0-THF, creatine phosphate and magnesium acetate in three different concentration sets and monitored the yield of active TC-Fluc at regular intervals thereafter by luciferase assay. While TC-Fluc synthesis in the control reaction without feeding stopped at 90 min, the replenished reactions continued for an additional 30 min. More importantly a ~2-fold increase in the maximum yield of active TC-Fluc compared to the control reaction, was reached at 120 min when 0.3 mM AA, 27 0D_260_/ml tRNAs, 0.1 mM DTT, 10 μg/ml 10-CH0-THF, 20 mM creatine phosphate and 2 mM magnesium acetate were fed (Supplementary Figure 4).

### Supplementing PURE system with missing protein factors isolated from *in vivo*

Some additional beneficiary factors present in *vivo* and in cell lysates but not in the PURE system also contribute to the sustainability of protein synthesis. Structural and functional studies suggested that EF-P promotes formation of the first peptide bond during proteins synthesis ^39^. More recent studies showed that EF-P could also act during elongation by facilitating the synthesis of proteins containing stretches of consecutive proline residues^31, 32^. To test if EF-P would improve translation efficiencies also on templates without any poly-proline stretches, we supplemented EF-P to TC-Fluc synthesis in the PURE system. Indeed, 4 μM EF-P (optimized by a pilot concentration gradient experiment, Supplementary Figure 5), increased the yield of the functional TC-Fluc translation-measured by luciferase – as much as ~2-fold (Figure 4A). Interestingly, the reaction with EF-P produced, first, significantly less active TC-Fluc before 30 min, about equal amount of active TC-Fluc at 30 min, and more active TC-Fluc after 30 min than the reaction without EF-P, eventually resulting in ~2-fold increase in activity by 120 min (Figure 4A). Paradoxically, the total yield of TC-Fluc determined by TC-tag labeling assay was lower in the presence of EF-P at all time points compared to the with no EF-P (Figure 4B). Although EF-P slows down translation by either slowing initiation or elongation rates, it is conceivable that a slower translation rate allows nascent peptide to fold properly and efficiently, therefore resulting in a larger fraction of active product^40^. EF4 recognizes ribosomes after a defective translocation reaction and induces a back-translocation, thus giving EF-G a second chance to translocate the tRNAs correctly^41^. EF4 boosts protein synthesis in living cells^33^ and the cell extract system even with elevated Mg^2+^ concentrations^41^ as well as in the classical PURE system^25^. Consistent with previous results, EF4 rescued high Mg^2+^ caused stress in a more defined system such as PURE (Supplementary Figure 6): EF4 (maximally with a EF4:ribosome molar ratio of 0.7:1) doubled the translation efficiency of active Fluc produced in the presence of high Mg^2+^ (+7 mM) stress (Supplementary Figure 6A). Similarly, the total amount of HaloTag AcGFP (HT-AcGFP) produced in the PURE system significantly decreased when magnesium acetate concentration increased by 7mM. Adding EF4 with a EF4:ribosome molar ratio of 0.7:1 increased not only the yield of total HT-AcGFP but also the amount of partial translated product (Supplementary Figure 6B).

**Figure 4.**
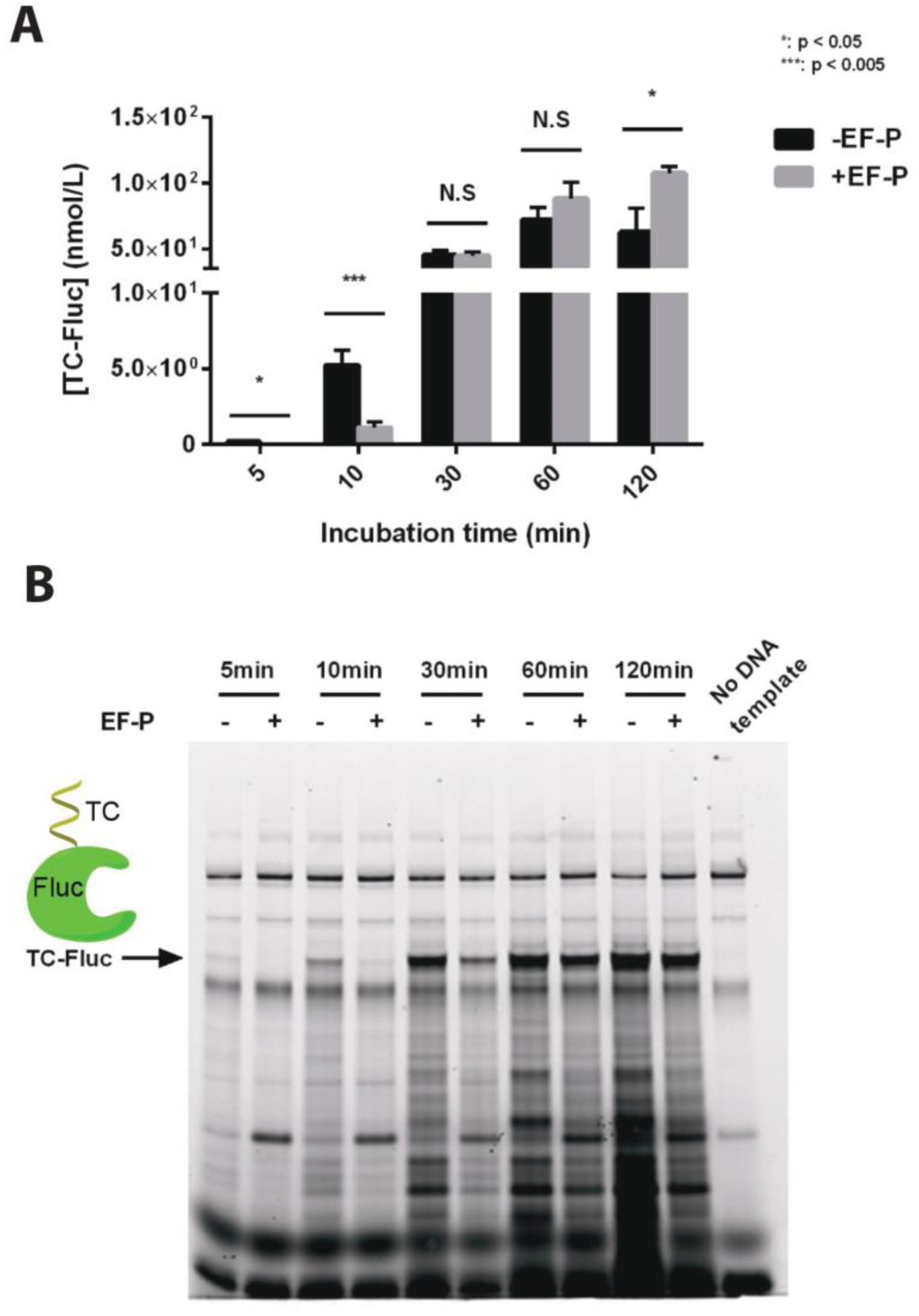
Supplementing the PURE system with EF-P. **A.** Time course study of active TC-Fluc production in the PURE system with or without 4 μM EF-P. TC-Fluc activity was measured after incubation at 37°C for the indicated time lengths. Values represent averages and error bars are ± standard deviations, with n=3. (*: p < 0.05; ***: p < 0.005; N.S: not significant) **B.** Time course study of total TC-Fluc produced in the PURE system with or without 4 μM EF-P. PURE system reaction aliquots were harvested at different time points, were incubated with FIAsH-EDT_2_, and separated by SDS-PAGE. Gel was scanned by a typhoon scanner with filter set (508nmEx/528nmEm). Arrow indicates full length TC-Fluc.

In addition to the tmRNA-dependent *trans-translation* system, two other ribosome rescue systems YaeJ and ArfA/RF2 can release stalled ribosomes from mRNAs lacking in-frame stop codons. However, previous studies only showed YaeJ and ArfA/RF2 performed this function when a short peptide of 7 amino acids was synthesized^28^ and role of these two systems on long peptides is still unknown. To examine the activities of YaeJ and ArfA/RF2 systems during the translation of a full length protein in the PURE system, we tested the addition of YaeJ and ArfA (RF2 already present in the PURE system) with three DNA templates: linear TC-Fluc with a stop codon; linear TC-Fluc lacking a stop codon; circular pIVEX 2.3d-TC-Fluc. As we expected, the addition of ArfA or YaeJ didn’t have any effect on the yield of TC-Fluc when linear TC-Fluc template with a stop codon and circular template pIVEX 2.3d-TC-Fluc were used. However, only ArfA improved TC-Fluc yield when linear TC-Fluc template with no stop codon was used while neither YaeJ (Figure 5A-B) nor PrfH (Supplementary Figure 7) showed any effect. Significantly, higher amount of TC-Fluc was produced with the circular template than the linear templates (Figure 5A-B). Quantifying fully and partially translated TC-Fluc with the TC tag showed that a larger fraction of ribosomes stalled on the mRNA transcribed from the linear templates than the circular one when translation was nearly accomplished (Supplementary Figure 8). This suggests that the 3’ sequence after stop codon encoding T7 terminator in the circular template helped mRNA form a secondary structure alleviating ribosome stalling, therefore resulting in a higher yield. Similar amount of TC-Fluc was produced when linear TC-Fluc with and without stop codon were used (Figure 5A-B), indicating stop codon was not the main limiting factor of ribosome recycling in this case.

**Figure 5.**
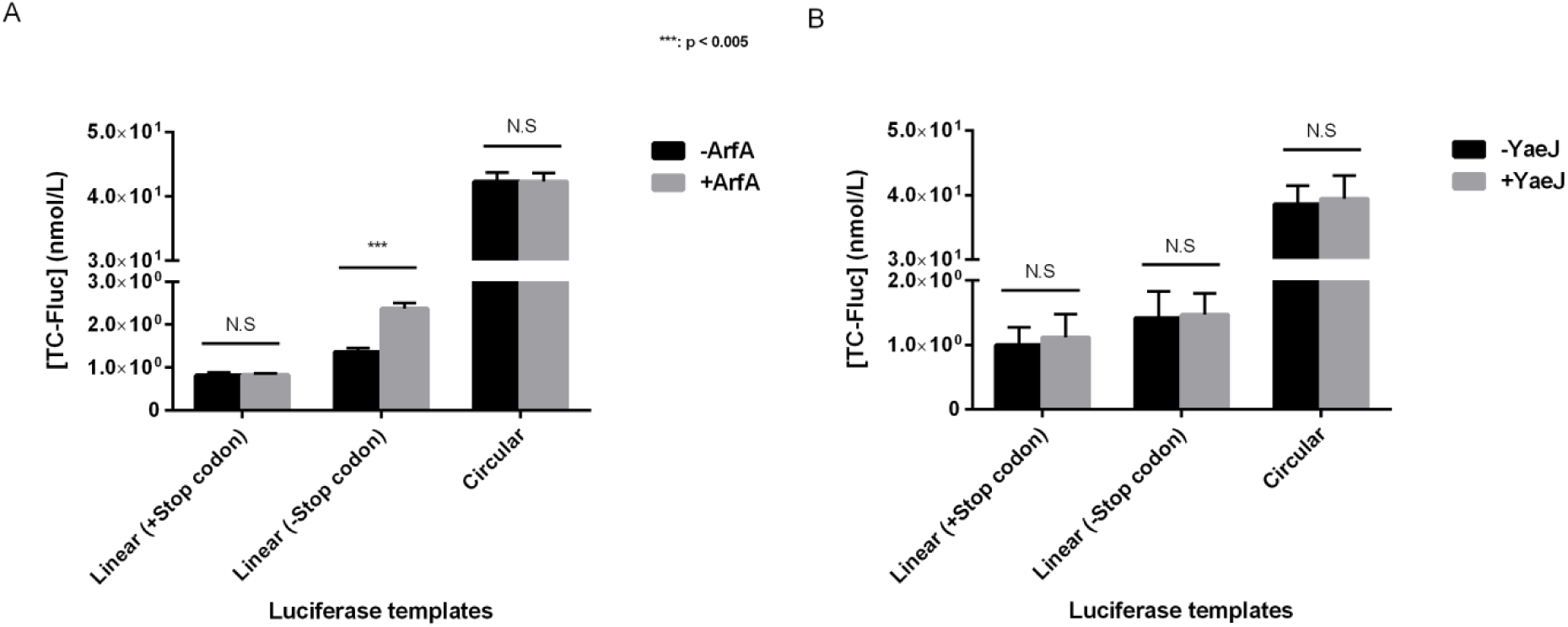
Supplementing the PURE system with ribosome rescuing systems. **A.** Effect of ArfA presence to the PURE system TC-Fluc synthesis. PURE system was assembled with different templates encoding TC-Fluc as indicated, with or without 8 μM ArfA protein. TC-Fluc activity was measured after 2 h incubation at 37°C. Values represent averages and error bars are ± standard deviations, with n=3. (***: p < 0.005; N.S: not significant) **B.** Effect of YaeJ presence to the PURE system TC-Fluc synthesis. PURE system was assembled with different templates encoding TC-Fluc as indicated, with or without 6 μM YaeJ protein. TC-Fluc activity was measured after 2 h incubation at 37°C. Values represent averages and error bars are ± standard deviations, with n=3. (N.S: not significant)

## DISCUSSION

CFPS systems have already been utilized for a number of applications such as directed evolution of proteins^5^, site-specific incorporation of non-standard amino acids^3, 4^ and has the potential for studying individual components of protein translation^25^. Toward building a self-replicating system, the utilization of already existing biological machineries promise to be a sensible strategy. A CFPS environment that consists of the optimized concentrations of proteins, energy source, monomer substrates, cofactors and DNA with desired genes, in theory, could constitute such a minimal, self-replicating dynamic system^25, 42^. Previous models propose that ~41,673 peptide bonds per ribosomes need to be synthesized in order to produce all the components of a self-replicating system^42^. For such a daunting task, beyond other cell-extract dependent CFPSs^43^, we approached PURE system as the ideal starting point^25^ because PURE allows controlled access to each of these components/conditions without being limited by contaminants such as endo/exo nucleases. However, even though classical PURE system contains wild-type ribosomes, it is still ~1-2 orders of magnitude less efficient than the predicted efficiency for a self-replicating system^44^. In this work, we tend to dissect the limiting factors of PURE protein synthesis by looking into individual components such as small molecule substrates and missing function modules involved in proteins synthesis.

We first made a side-by-side comparison of PURE reaction with S30 system in terms of transcription and translation. Although PURE resulted total protein yields up to ~4 times more than S30; the formation of the functional protein by S30 was up to ~3-fold higher (Figure 1C-D), which suggests presence of additional factors in S30 perhaps improving translation fidelity and facilitating proper protein folding and/or post-translational modifications. Furthermore, while a comparison of the expression profiles of reporter proteins in these two systems clearly indicated non-specific RNA degradation in S30 (probably due to nuclease contamination) that was absent in PURE (Figure 1A-B). However, PURE too showed a substantial, characteristic smear of partially translated reporter proteins suggestive of premature protein translation termination due to the lack of rescue factors and mRNA chaperones that appear to be present in S30 (Figure 1C-D). We tested the expression of different HaloTag reporter proteins with sizes ranging 90 kD – 220 kD in the PURE and S30 systems. We found the yields of protein translation tend to decline as protein size increased and both of the two systems showed low ribosome recycling efficiency (Figure 2).

The lower yields of functional proteins with classical PURE are likely a result of low translation efficiency of the PURE ribosome and misfolding of the nascent polypeptides, suggesting that PURE might be lacking a number of protein components such as chaperones^25, 45^, translation facilitating factors and efficient recycling modules resulting in the depletion of energy and monomers or accumulation of inhibitory byproducts.

Expression profile of these proteins in PURE over time confirmed that the new protein synthesis stopped ~2 h (Supplementary Figure 1). Interestingly, individual fed-batch replenishment of the PURE components at later time points (~75 min), in general, resulted in higher reaction yields within a given time point prior to the stop of translation (90 min) (Figure 3) suggesting that this treatment can boast the rate of protein translation. Indeed, replenishment with a combination of these components at later points not only extended the reaction duration to about ~120 min with a ~2-fold yield increase, but also boosted the rate of functional protein translation (Supplementary Figure 4). This increase in yields and rates could be explained by countering substrate/energy depletion and/or time-dependent degradation of some of the components (e.g. DTT).

When the system was replenished with the said components in earlier time points, this boost was not seen, probably because of a variety of inhibition modes by the increased substrate concentrations. For example, higher NTP concentrations at earlier time points were inhibitory (Figure 3G) likely because of the chelation of the necessary Mg^2+^ cations by excess NTPs^46^. Indeed, supplying the system with magnesium acetate had a beneficiary effect on yields at any time-point (Figure 3F), probably addressing Mg^2+^ cation chelation by the accumulating inorganic phosphate created by the creatine phosphate shuttle.

The PURE system still lacks some crucial recycling components and chaperone systems relaxing mRNA secondary structures. A close inspection of other factors known to affect protein translation^47^ but absent in the classical PURE system, brought up EF-P as a candidate which is known to reverse protein translation stalling on poly-proline stretches^31, 32^. Indeed, EF-P increased the yield of functional protein 2-fold at the latest time points (~120 min) even though the total protein yields were lower at all time-points (Figure 4). This is consistent with EF-P being known to slow down protein translation at the stage of initiation^48^ and/or elongation, which might facilitate a more efficient folding of the nascent peptide into a functional protein as well as by lowering premature protein termination events. EF4, just like in the cell extract system^41^ and living cells^33^, boosts protein translation not only in the classical PURE system^25^, but also in the PURE system with an elevated Mg^2+^ concentration (Supplementary Figure 6). Unlike translating short peptides, YaeJ or PrfH didn’t help on ribosome recycling when translating the large protein TC-Fluc with a linear DNA template with no stop codon, although ArfA still performed the function in this case (Figure 5 and Supplementary Figure 7). It indicates other factors may be needed by YaeJ or PrfH to recycle ribosomes when translating large proteins. Besides, our observation also suggests mRNA secondary structure is another factor leading to ribosomes stalling and therefore influences total product yield. Adding mRNA chaperones could possibly increase system productivity by reducing ribosome stalling.

Toward a CFPS-based self-replication system, utilization of PURE system is particularly attractive due to its modularity. Achieving the theoretical translation efficiencies required for a self-replicated system with PURE will likely require inclusion of additional proteins/modules that may be inherently crucial for protein translation in cells. Indeed, here we show that this optimization process can involve substrate replenishment, by-product recycling and addition of new protein factors. At the same time, this process can also provide us with new insights and facilitate us to test hypotheses about potentially important but poorly characterized players in protein translation, as it did here about EF-P, EF4, ArfA, YaeJ and PrfH.

## MATERIALS AND METHODS

### Media, chemicals, reagents and device

Unless specified, all chemicals were obtained from Sigma-Aldrich. For protein purification, liquid cultures of strains were grown in either SB media (24 g/L tryptone, 12 g/L yeast extract, 5 g/L glucose, 2 g/L NaH_2_PO_4_, 16.4 g/L K_2_HPO_4_-3H_2_O, 4 mL/L glycerol) or 2YT media (16 g/L Tryptone, 10 g/L yeast extract, 5 g/L NaCl). Purified protein concentrations were determined by standard Bradford assay (Bio-Rad). Phusion High-Fidelity PCR Master Mix, restriction enzymes, Quick Ligation Kit and PURE system were purchased from NEB. RTS 100 *E.coli* HY Kit (S30 system) was purchased from 5PRIME. The Pierce 3.5K MWCO Microdialysis Plate was obtained from Thermo Fisher Scientific (Rockford, IL).

### Molecular cloning and protein purification

The Fluc gene and TC-Fluc was cloned into the NcoI and XhoI restriction sites of pIVEX-2.3d (5 PRIME). Linear templates encoding Fluc or TC-Fluc starting from T7 promoter and ending right before or at stop codon were generated by PCR for further use in PURE system reactions. Cloning of plasmids encoding HT-Ch-TolA, HT-EntE-Ch-TolA and HT-EntF-Ch-TolA was described in ^25^. The vector pLJ for expressing N-terminal HaloTagged AcGFP was derived from pFN18K (Promega). Specifically AcGFP was inserted after the sequences of an N-terminal HaloTag domain. PIVEX 2.3d-Fluc, pIVEX 2.3d-TC-Fluc, pLJ, linear Fluc and TC-Fluc templates were purified with phenol: chloroform: isoamyl alcohol extraction and ethanol precipitation before being used for in vitro transcription and translation.

EF4 was cloned, overexpressed and purified as described in ^25^. The genes encoding YaeJ and ArfA were cloned from *E. coli* MG1655 genomic DNA (ATCC) with a C-terminal histag and inserted into pET-24b (Merck Millipore) via NdeI and XhoI. The 12 C-terminal amino acid residues of ArfA were deleted because these residues lead to poor expression and their deletion does not affect ArfA activity ^28, 49^. PET-24b YaeJ and ArfA were transformed to BL21 (DE3) strain (NEB). Cells were grown in LB with 50 μg/mL kanamycin at 37 °C, induced with 1 mM IPTG when OD600 reached 0.6, and further incubated at 37 °C to an A600 of 1.4–1.8. Cells were recovered by centrifugation and pellets were washed by binding buffer (20 mM sodium phosphate, 0.5 M NaCl, 40 mM imidazole, 6 mM β-ME pH 7.4) 3 times and lysed by BugBuster Protein Extraction Reagent (Novagen) with 100 μL Halt-protease inhibitor (Thermo-Fisher), 6 mM β-ME in binding buffer. Cell lysates were centrifuged at 15,000g for 25 min. The supernatant was applied to GE histrap-Gravity column and washed with binding buffer. Histagged enzymes were eluted by elution buffer (20 mM sodium phosphate, 0.5 M NaCl, 500 mM imidazole, 6 mM β-ME pH 7.4), pooled, concentrated with Amicon-Ultra-4 concentrator with 3K MWCO, dialyzed to a buffer containing 50 mM Tris–HCl pH 7.6, 100 mM KCl and 1 mM DTT for 6 h twice and then a storage buffer containing 50 mM Tris–HCl pH 7.6, 100 mM KCl, 1 mM DTT and 30% glycerol. Purified YaeJ and ArfA were stored at −80 °C.

For EF-P preparation, plasmid pST39-his-efp-yjeA-yheK, which encoding EF-P and its modification enzymes was kindly provided by Dr. Park Myung Hee at NIH. 0verexpression and purification of EF-P was performed by following the protocol from ^50^. Specifically, the plasmid was transformed into BL21 (DE3) pLysS strain (Agilent) and grown in 5 mL of 2YT media with 100 μg/mL of Carbenicillin overnight. 250 mL media were inoculated with the overnight cultures and protein expression was induced with 1 mM of IPTG at OD600 around 0.6-0.8. The cells were incubated at shaker at 20°C for 16 h and harvest by centrifuge. Cell pellet was then resuspended in Buffer A (50 mM Hepes-K0H pH 7.6, 1 M NH_4_Cl, 10 mM MgCl_2_ and 7 mM β-ME) and lysed with french pressure cell. The protein purification was carried out with ÄKTAprime (GE Healthcare) equipped with 5 mL HisTrap™ HP column (GE Healthcare). EF-P was eluted into Buffer A plus 200 mM Imidazole and dialyzed into buffer C (50 mM Hepes-K0H pH 7.6, 100 mM KCl, 10 mM MgCl_2_ and 7 mM β-ME). EF-P was concentrated to 0.5 mL and stored in −80°C upon adding one volume of 80 % glycerol.

PrfH gene was cloned from *E. coli* MG1655 genome into pGEX-6p-1 vector and transformed into BL21 (DE3) pLysS strain. A single colony was inoculated and grown in 5 mL of 2YT media with 100 μg/mL of Carbenicillin overnight. 50 mL media were inoculated with the overnight cultures and protein expression was induced with 1 mM of IPTG at room temperature for 3 h before harvesting via centrifugation. Protein purification was performed with Glutathione Sepharose 4B according to manufacturer instruction, and cleaved off the sepharose beads with PreScission Protease at 4°C in cleavage buffer containing 50 mM Tris pH 8.0, 150 mM NaCl, 1 mM EDTA, 1 mM DTT, 1% Triton X-100. PreScission Protease cleavage generated a single protein product corresponding to PrfH (with no affinity tags) based on Coomassie Blue staining.

### Total RNA extraction and electrophoresis

PURE system and S30 system (RTS 100 *E.coli* HY Kit) reactions were set up according to manufacturer's instructions. Total RNA was extracted using phenol-chloroform extraction at the end of the reactions. 1μg of each RNA sample was denatured and subjected to electrophoresis in 1% agarose gel, followed by visualization with SYBR Green dye (Life Technologies).

### Luciferase assay

Relative luminescence units were measured by a microplate reader. The amount of functional TC-Fluc produced was measured by converting relative luminescence units to the actual amount of TC-Fluc via a premeasured standard curve.

### Labelling of in vitro translated proteins

To further examine full-length and incomplete products of *in vitro* translation in PURE system, HT-Ch-TolA, HT-EntE-Ch-TolA, HT-EntF-Ch-TolA and HT-AcGFP were *in vitro* translated and labelled with a fluorescent TMR reporter (Promega); TC-Fluc and TC-Fluc-chis were *in vitro* translated and labelled with FlAsH-EDT_2_ biarsenical labeling reagent (Life Technologies). Specifically, reactions expressing HT tagged proteins were diluted by 10 times with PBS and incubated with 5 μM TMR ligand at room temperature for 30 min prior to SDS-PAGE analysis. Gels were scanned at 580 nm (excitation at 532 nm) by Typhoon Trio Imager (GE Healthcare) and a fluorescence gel image was analyzed with ImageQuant TL software. For labeling of TC-Fluc and TC-Fluc-chis, PURE system and S30 system reactions were first diluted 1:10 in PBS and mixed with 1.25 μM FlAsH-EDT_2_ biarsenical labeling reagent (diluted in PBS). The mixture was incubated at 25 °C for 30min, and then subjected to SDS-PAGE analysis. Fluorescent signal was obtained with Typhoon Trio Imager (GE Healthcare) at 528 nm (excitation at 508 nm).

### PURE fed-batch reactions

PURE batch reactions (10 μL) synthesizing TC-Fluc were performed following the manufacture's instructions. The volumes of water or substrates (individual and in combination, prepared to proper stock concentrations to give desired replenishment) added to PURE were set to 1 μL to avoid dilution effect. We first set up a series of 10 μL PURE reactions and added 1 μL of water at time 0 min, 15 min, 30 min, 45 min, 60 min or 75 min to the reactions. TC-Fluc was quantified at 90 min by luciferase assay to make sure these testing reactions produced similar amount of TC-Fluc (data not shown). Therefore adding 1 μL water at time 0 min can be used as control for the fed-batch tests of substrates. An equivalent volume of feed substrates were added directly into the reactions at given time points to replenish the desired substrate at concentrations specified in the text.

### Inorganic phosphate quantitative assay

Quantitative analysis of inorganic phosphate was performed using the EnzChek Phosphate Assay Kit (Life Technologies). PURE samples were first quenched by frozen on liquid nitrogen and later thawed and assayed. To measure phosphate concentration, samples were diluted 200-fold with MQ water and compared to a linear phosphate standard ranging from 2 to 150 μM. Absorbance was read at 360 nm following the manufacture's instructions.

### PURE transcription and translation with translation factors

5 μL reactions were carried out following the PURE system manual with 2 μL solution A, 1.5 μL solution B, 0.8 U/μL SUPERase IN RNase Inhibitor (Life Technologies), 50 ng circular or linear DNA template, and proper amount of supplement factors as indicated in the text. Reactions were incubated at 37 °C for 2 h unless otherwise stated. For analyzing the effect of EF-P on TC-Fluc activity at different time points, samples for prior time points were stored at 4°C till the last time point was reached, and TC-Fluc activity and/or fluorescent labeling was analyzed for all samples together. Fluc activity was measured by Promega Luciferase Assay System kit. Chemical luminescence in relative luminescent units (RLUs) was measured by a microplate reader (SpectraMax M5, Molecular Devices).

## ACKNOWLEDGEMENTS

We thank Dr. Anthony Forster for comments and suggestions. This work was funded by programs from the Department of Energy (Genomes to Life Center) [Grant #DE-FG02-02ER63445].

## AUTHOR CONTRIBUTIONS

JL designed the study. JL, CZ, PH, TL and EFB performed the experiments. JL and CZ analyzed the data. JL and EK wrote the manuscript. All authors reviewed the manuscript. GMC performed a supervision role.

## DATA AVAILABILITY STATEMENT

All data underlying the findings described in the manuscript are fully available without restriction.

